# From neural network to psychophysics of time: Exploring emergent properties of RNNs using novel Hamiltonian formalism

**DOI:** 10.1101/125849

**Authors:** Rakesh Sengupta, Anindya Pattanayak, Raju Surampudi Bapi

## Abstract

The stability analysis of dynamical neural network systems generally follows the route of finding a suitable Liapunov function after the fashion Hopfield’s famous paper on content addressable memory network or by finding conditions that make divergent solutions impossible. For the current work we focused on biological recurrent neural networks (bRNNs) that require transient external inputs (Cohen-Grossberg networks). In the current work we have proposed a general method to construct Liapunov functions for recurrent neural network with the help of a physically meaningful Hamiltonian function. This construct allows us to explore the emergent properties of the recurrent network (e.g., parameter configuration needed for winner-take-all competition in a leaky accumulator design) beyond that available in standard stability analysis, while also comparing well with standard stability analysis (ordinary differential equation approach) as a special case of the general stability constraint derived from the Hamiltonian formulation. We also show that the Cohen-Grossberg Liapunov function can be derived naturally from the Hamiltonian formalism. A strength of the construct comes from its usability as a predictor for behavior in psychophysical experiments involving numerosity and temporal duration judgements.

## I. Introduction

Recurrent neural network (RNN) consists of interconnected neurons that have some feedback loops built-in between the nodes. The feedback can come from the same or different node at each time point of the network’s evolution and the behavior of the network resembles a nonlinear dynamical systems. These networks have been used for constructing neural models for memory [1], decision making [2], visual sense of numbers [3] to name a few. Some dynamic vision algorithms include recurrent neurons as building blocks for their usefulness in integrating and spreading local and global influences across the network [4]. The biological RNNs (bRNN) are mostly imagined as a single layer of neurons.

The on-center off-surround kind of recurrent networks have been very popular in the literature involving short-term memory, decision making, contour enhancement, pattern recognition, and several other problems because of their versatility and ability to produce self-organized outputs based on underlying non-linear mathematical properties of the network. One of the standard approaches towards the stability of these networks has always been to find a suitable Liapunov function or finding the condition for which network trajectories do not diverge [5], [6]. A detailed review of Liapunov and other stability approaches can be found in [7]. In the current paper we show a novel intuitive general purpose method for constructing a Liapunov function for recurrent networks in general and show several specific cases to show the effectiveness of the formalism. We also compare the stability criterion obtained from Hamiltonian formalism with ordinary differential equation approach. Towards the end of the paper we show how such general purpose construct can be used to generate predictions in actual biological systems.

## II. Hamiltonian Function For General Recurrent Network

### A. Cohen-Grossberg Liapunov function derived from Hamiltonian formalism

We begin by considering a single layer of fully connected recurrent neural nodes. The activation of *i*−th node is given by *x*_*i*_. If we consider a general recurrent shunting networks with dynamics given by^1^

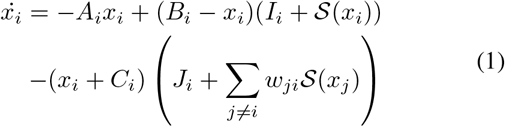
where *I* and *J* are excitatory and inhibitory inputs to the node *i* and 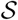 is a sigmoid function. *B* and −*C* are constants determining the upper and lower bound for the network activation respectively. We can transform Eq. 1 with symple variable change (*y_i_* = *x_i_* + *C_i_*) to the form

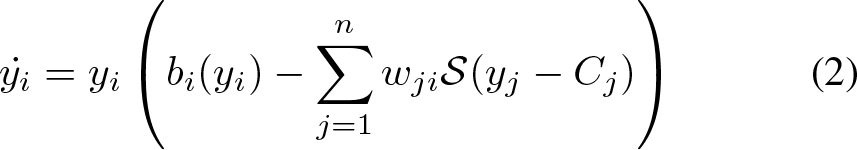

where,

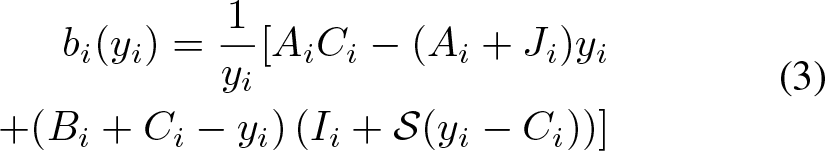

As *x_i_* is the activation of a particular node *i* of a recurrent network with *n* nodes, the general time evolution of all Cohen-Grossberg systems can be written as

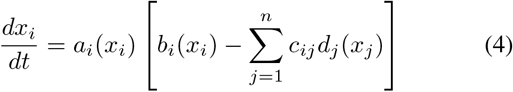
where *c_ij_* are symmetric coefficient. The above formulation is in fact quite general and the equation can be used for both additive and shunting model networks, continuous-time McCulloch-Pitts model, Boltzmann machines, mean field model among others^2^.

The global Liapunov function used by Cohen and Grossberg is given by [5], [6]

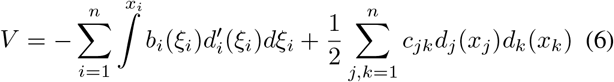

The steady-state or equilibrium solution in the time evolution of the network implies that the activations of the nodes will be unable to influence each other in the long-term, i.e., all the local instabilities will decay. It allows us to consider the set {*x*_*i*_} as a set of generalized co-ordinates describing the state of the network. It follows then that we can use the Hamiltonian principle from classical mechanics and say

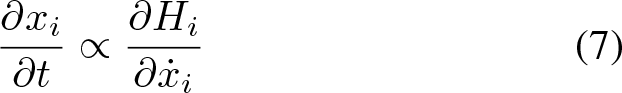

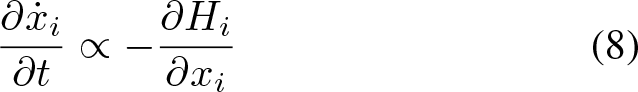

where 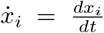. If the proportionality constants in Eq. 7 and 8 are unequal (the equality leading to a trivial case), the Hamiltonian for a particular node *i* will follow

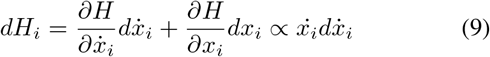

In the above we have used the identity 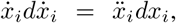, anticipating the final derivation.

Using Eq. 4, we have

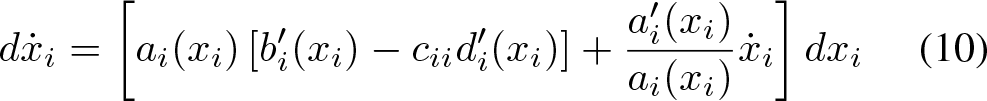

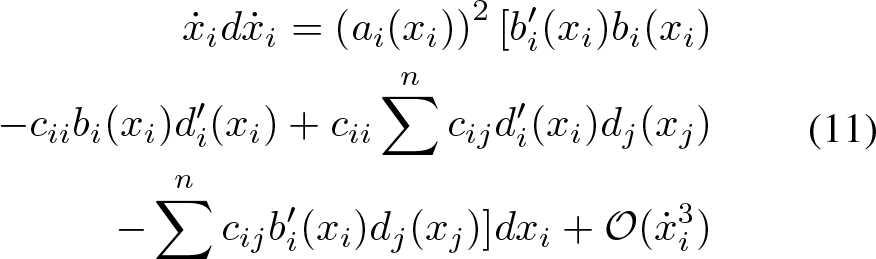

where 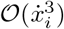 represents terms that are of the order to 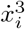.

Near equilibrium (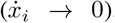), we can safely ignore the terms 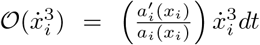 and also 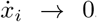 we have *b*_*i*_(*x*_*i*_ → Σ *c*_*ij*_ *d*_*j*_ (*x_j_*) and thus the first and last terms in the sum within parenthesis cancel each other. Ignoring multiplicative coefficients, we can say that near equilibrium,

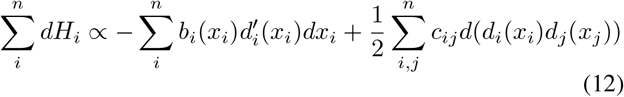

and thus the full Hamiltonian for the system can be written as (with some changes in the dummy indices),

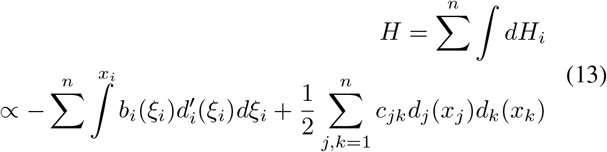

Thus we can show that the Cohen-Grossberg Liapunov function is a special case of and can be derived from general Hamiltonian formalism. This has a far reaching consequence for neural networks. As Eq. 4 is applicable for a large variety of neural networks, we can now derive energy functions based on the first principles rather than inspired guesses.

In the next section we will show certain special cases of the general formalism and how stability criteria could be found for several different kinds of recurrent networks. We also compare the stability criterion from the Hamiltonian with stability criterion derived from divergence tests.

### B. Coupled oscillatory brain network from the Hamiltonian formalism

Another interesting contribution of the Hamiltonian formalism is its application towards oscillatory neural theories. From Eq. 4 we can say that

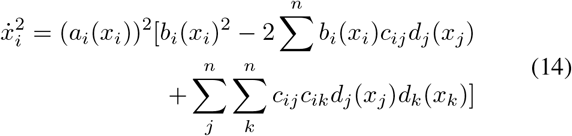

Since near equilibrium, we have *b_i_*(*x_i_*) → Σ *c_ij_ d_j_* (*x_j_*), we can write,

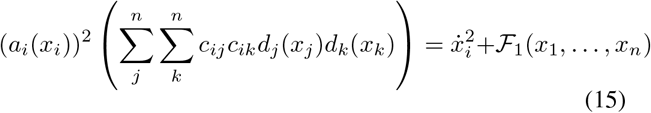

If 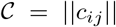 is a symmetric matrix (*c_ij_* = *c_ji_*), if the following property is satisfied

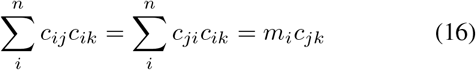

then *m_i_* are the diagonal elements of the diagonal matrix *D* which satisfies the following matrix relation

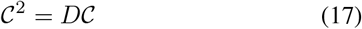

So using the multiplicative coefficients in Eq. 11, and using Eq. 15, 16 and 13, we can write the full Hamiltonian as the following

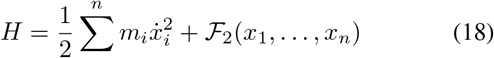
where *m_i_* are proportionality constants. Interestingly, the form of the Hamiltonian given in Eq: 18 is the Hamiltonian for multiple coupled nonlinear oscillators with equation of motion of the form [9]

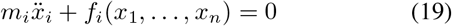

It is interesting that for the Hamiltonian to generate meaningful oscillatory solution, we need transient input to the system as well as a period where the network will be allowed to settle or stabilize. Thus for oscillatory dynamics to be biologically feasible in the brain we need a recurrent layer to get transient input from a feed-forward network. The above formulation gives very important insights and constraints of deriving oscillatory models of the brain function.

## III. SPECIAL CASES OF HAMILTONIAN FORMALISM

A recurrent shunting network with the range [−*D, B*] follows the dynamics

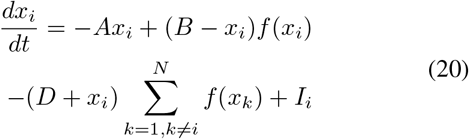

A general additive recurrent network is given by

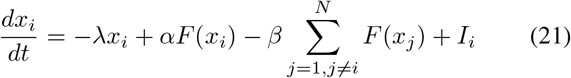

For the additive network we assume a decay constant of *λ*.

### A. Additive Recurrent Network with Slower than Linear Activation Function

Let us assume that in Eq. 21,

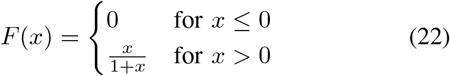

The network should reach steady state activity when the external input is taken away. If we disregard noise, at steady state, i.e. when, 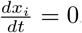,

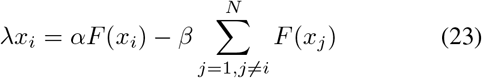

As the equation is symmetric under permutation of units, the system should have symmetric solutions characterized by number of active units *n*, and their activation *x*(*n*), all other units having 0 activation.

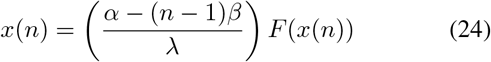

Using Eq. 22, we get

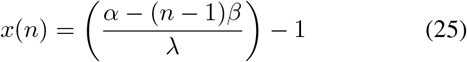

Noise can bring in additional fluctuation that can destabilize the solution for a pair of active modes (with equal activation according to Eq. 25), unless the the difference of activations between the said nodes Δ*x* = *x_i_ − x_j_* decays. Using Eq. 21 & 25 we get

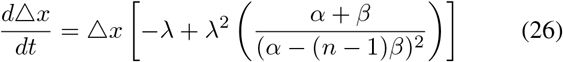

Thus the fluctuation decays only if 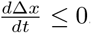, i.e.,

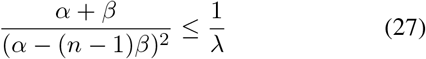

As we can see that the decay parameter, excitation parameter and inhibition parameter are not completely independent for stable solutions. For the present purposes we use *λ* = 1 (we will keep using this value for the rest of the paper for simplicity).

From Eq. 21 and 22 we have

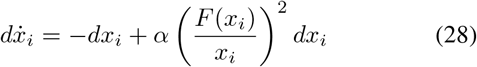

It is easy to show that from the definition given in 9,

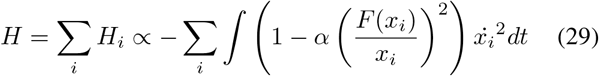

From Eq. 25 we can substitute terms in steady state to get

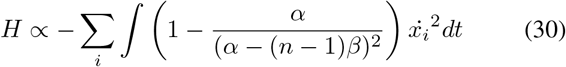

Now if *dH <* 0 and thus a monotonically decreasing Liapunov type function in absence of external input, we have the stability condition as

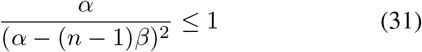

Comparing this to Eq. 27, we can see that the conditions derived from the energy value is slightly different and diverges greatly for higher *β*. This is due to the fact that Eq. 27 excludes winner-take-all mechanisms operating at higher inhibition, whereas the energy function does not. And it is evident that for all *β*,

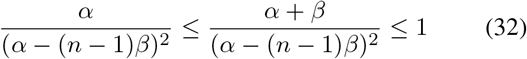

and thus the energy function is a very suitable candidate for the network as it is in line with the stability analysis derived from the dynamics of the network.

### B. Shunting Recurrent Network with Constant Activation Function

Here we assume in Eq. 20,

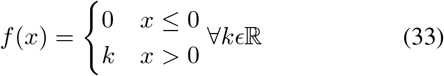

A search for a symmetric solution at steady state leads to steady state activation value

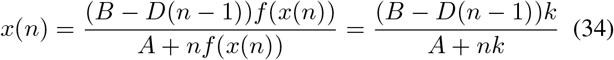

The stability criterion is determined considering the decay of Δ*x* = *x_i_ − x_j_* and is calculated to the following condition

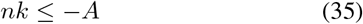

Constructing the energy function using

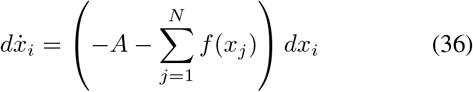

and Eq. 9, we have
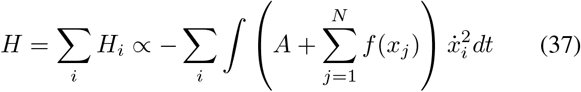

As 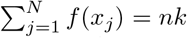near steady state, we have the same criterion for stability from the energy function as Eq. 35.

### C. Shunting Network with Linear Activation Function

The activation function for such networks is given by

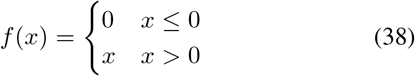

Using the above activation function in Eq. 20 we have at steady state condition,

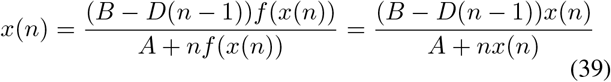

This leads to the non-trivial solution

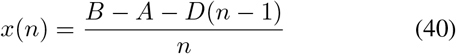

Looking at the decay of Δ*x* = *x_i_ − x_j_* leads to the stability criterion,

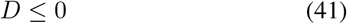

The Hamiltonian constructed using

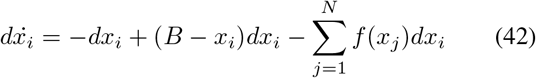
and Eq. 9 leads to,
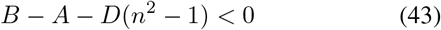
which upon substituting *B − A* = *nx*(*n*) + *D*(*n −* 1) (from Eq.40) yields *nx*(*n*) *− nD*(*n −* 1) *<* 0. For the situation where total network activation *nx*(*n*) *≥* 0, we have *D ≥* 0 for *n >* 1. In fact this leads to a better stability criterion combining the stability criterion given in Eq. 41 with the condition *D ≥* 0 obtained from the Hamiltonian
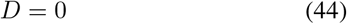

This is a sensible result as by definition of shunting networks *B* and *D* are positive real quantities.

### D. Shunting Network Using Reciprocal Activation Function

The activation function for such networks is given by

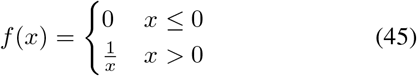

The steady state solution (assuming the positive root of a quadratic equation) is given by,

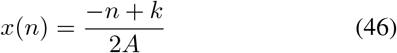
where 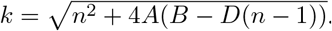

The stability criterion from decay condition is derived to be
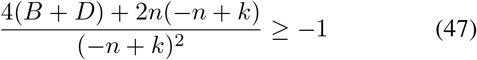

Constructing the energy function in the now familiar way, we get

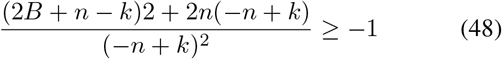

It can be shown that both the criteria given by Eq. 47 and 48 are same if *n − k* =2*D*.

## 4 Analytical Prediction of Winner-Take-All

In section III-A we have shown that there are constraints on the stability of the network response in an additive recurrent model, i.e., the condition that allows for the activation of two nodes to not diverge during the simulation. The constraint was given by

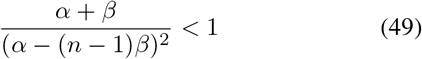
where *n* is the number of nodes active after the simulation is run. We also obtained another similar stability condition after consideration of the novel Hamiltonian formulation, mainly,

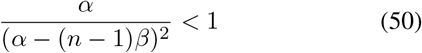

However, for a very important behavior of the network, the winner-take-all dynamics requires activation differences between nodes to be amplified until only one node emerges as winner. Now if *n* = 1, as in standard WTA interaction, Eq. 50 transforms into

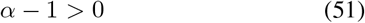

This gives the lower limit of *α* for WTA. The upper limit will be set by Eq. 49, i.e. when *n* = 2 reaches stability, i.e.,

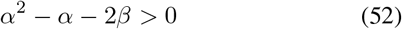

Thus WTA behavior will be supported by a range of *α* that satisfies the conditions *α ≥* 1 and *α*^2^ *− α −* ^2^*β ≤* 0. This is shown in Fig. 1. It shows how the *α* ranges should be calculated for different *β* values. For *β* = 0.25 the range is 1 *≤ α ≤* 1.37, for *β* = 0.3, 1 *≤ α ≤* 1.42, for *β* = 0.35, 1 *≤ α ≤* 1.47.

**Fig. 1.**
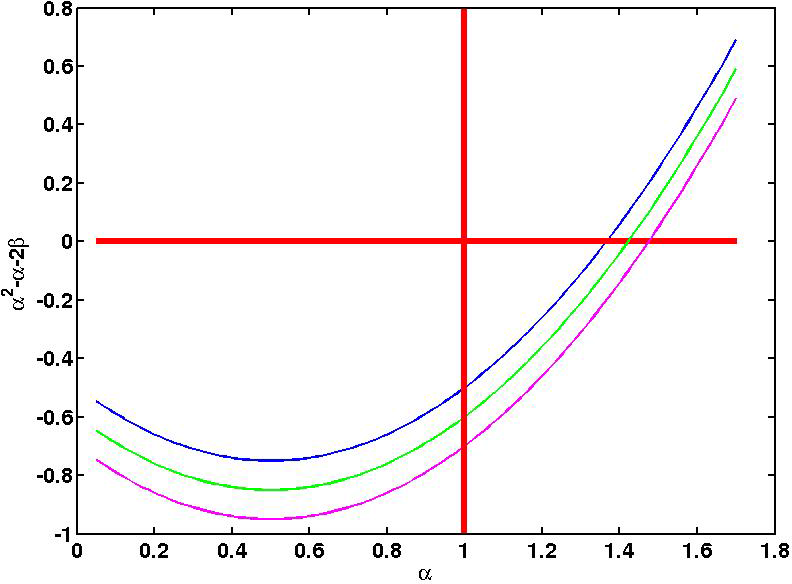
A plot of *α*^2^ *−α−* 2*β* for different *β* values. Blue line for *β* = 0.25, green for *β* = 0.3 and magenta for *β* = 0.35. In the bottom right quadrant of the plot bound by the lines *α*^2^ *− α −* 2*β* = 0 and *α* = 1 we get the *α* values desired for WTA interaction, mainly *α ≥* 1 and *α*^2^ *− α −* 2*β ≤* 0. For *β* = 0.25 the range is 1 *≤ α ≤* 1.37, for *β* = 0.3, 1 *≤ α ≤* 1.42, for *β* = 0.35, 1 *≤ α ≤* 1.47.

We also confirmed the parameters with actual simulation. We chose a network of 10 nodes and gave inputs to 2 nodes for each simulation. The input level was clamped at 0.3. The probability of WTA interaction was calculated as the fraction of time out of 1000 simulations, that only one node survives. We varied the *α* between 0.5 and 1.5. Both stimuli were presented for 255 time steps each and the total duration of simulation was 2500 time steps. Noise was sampled from normal distribution of mean 0 and standard deviation 0.1. The results are shown in Fig. 2. The analytic limits obtained in Fig. 1 are closely confirmed by the simulation as well. The specific parameter values for the simulation are given in Table I.

**Fig. 2.**
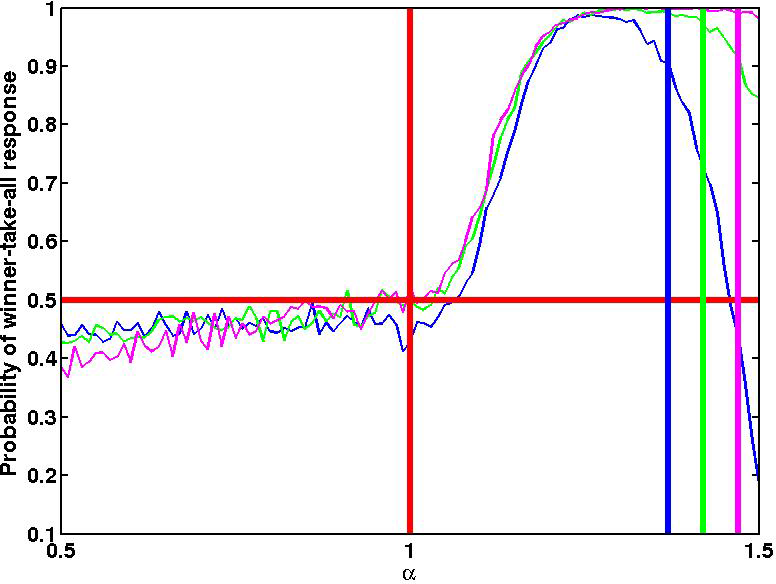
Probability of winner take all interaction plotted against *α* for three different *β* values 0.25 (blue line), 0.30 (green line) and 0.35 (magenta line). For *α >* 1, the WTA probability increases beyond chance level (0.5). The analytical limits on *α* obtained in Fig. 1 are shown in correspondingly colored lines.

## V. Applicationofwta in Temporal Oddball Experiments in Psychology

Subjective expansion of time (TSE) is the phenomenon where human beings perceive an oddball stimulus in a stream/ series of identical standard stimuli to have lasted longer in terms of percept duration [10]. There are several competing theories that have tried to explain this phenomenon. However, they can be roughly classified into - a) dedicated internal clock b) information based decision mechanism [11]–[13]. Here we have used a computational model to explore the hypothesis that recurrent on-center off-surround network [3] (with the hypothesis that oddball is more perceptually salient than standard stimuli) executing a winner-take-all decision can account for the oddball effect.

In the context of perceptual decision making, a winner-take-all mechanism is a prime candidate for a biologically plausible neuro-computational approach [14]. In case of temporal oddball paradigm, we observed that the experiment follows a two-alternative forced choice (2AFC) task. The 2AFC task led to the idea that time’s subjective expansion factor observed in temporal oddball paradigm might arise from a flexible decision boundary arising from a neural WTA mechanism.

The on-center off-surround architecture we have used here has been described in more detail in [3], [15]. Here we will just mention the salient points. The differential equation governing the time-evolution of the network of *N* nodes is given by

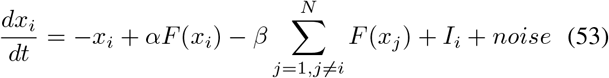

*x_i_*(*t*) is the activation of node *i* at time *t*. *I_i_* represents the intensity of external input (*∀i,* 0 *≤ I_i_ ≤* 1), it is zero if the stimulus is absent for a particular node at that time point. Input is only presented for a finite amount of time, typically much less than total time of simulation. *F* (*x*) is the activation function given by the formula,

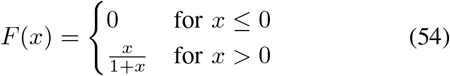

The network has a stable dynamics, i.e., the activated states at equilibrium do not diverge from the steady state dynamics if

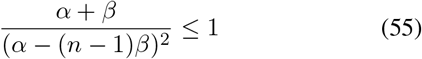

Considering a Liapunov function for energy for the same network we get another stability criterion

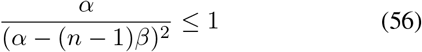

Considering these two we get a condition for winner-take-all (WTA) interaction for a range of *α* that satisfies the conditions *α ≥* 1 and *α*^2^ *− α −* 2*β ≤* 0. After some rigorous convergence tests we used the following modeling parameters In a standard oddball task for temporal judgements involve showing a participant a series of frequent standard stimuli followed by infrequent oddball stimulus. A few standard frequent stimuli might also be shown after oddball presentation. The participant makes a judgement whether the oddball stimulus was longer in duration than the standard or not. Our model assumes that a normalized input is given to two nodes of varying durations simultaneously as the duration judgements are post-event judgements in temporal oddball, it is not such an unreasonable assumption.

Now we ran 100 simulations for each duration pairs (30 vs 660 ms steps, 75 vs. 600 ms steps,… 1200 vs. 660 ms steps) for the two nodes and calculated the probability of response that one duration will be judged longer than the other by counting the fraction of times the node with the variable input is the winner of WTA interaction. In reality it is rare that both the units to be excited with the same level of input. It is more likely that the salience of the inputs differ. The standard stimuli is more frequent and thus prone to habituation and should receive lesser level input than the oddball stimuli. So the temporal oddball should correspond to the case where the standard interval gets a clamped input at level 0.2 and oddball interval gets clamped input at 0.3.

Now we calculated the values for the probability that oddball duration would be perceived as longer for each of the standard durations in the set (30:45:1200) (to be given input of 0.2). The oddball durations are also (30:45:1200) (to be given input 0.3). We calculated the points of subjective equality (PSE) through sigmoid fits and time’s subjective expansion by dividing the standard durations by corresponding PSE’s. The result is given in Fig. 3. The overall pattern of the results is strikingly close to TSE pattern observed in experimental results given in [10].

**Fig. 3.**
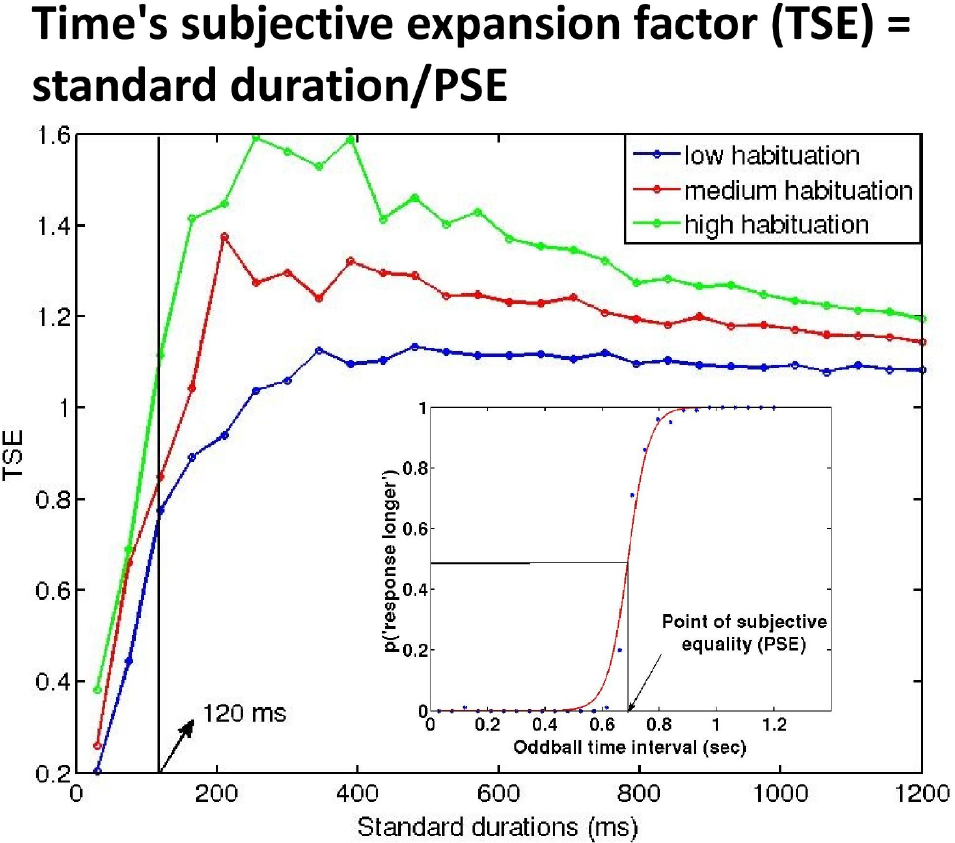
TSE values for different standard durations at different levels of oddball input. Blue line has oddball input at 0.3, red line has oddball input at 0.4, green line has oddball input at 0.5. The different levels were interpreted as arising from different levels of habituation, i.e., if the oddball appears in a stream of 6 standard stimuli, its salience would be less than that appearing in a stream of 11 stimuli, and thus the former will have lower stimulus level (see appendix). The inset shows an example psychophysical curve generated in order to calculate the point of subjective equality.

## VI. DISCUSSION

In the current work we have shown how analytical Hamiltonian formalism developed by the authors can derive Cohen-Grossberg Liapunov function for general single-layer shunting recurrent neural networks. This has a far reaching consequence the dynamics of such networks can be used to model a wide variety of neural networks including additive recurrent networks, continuous-time McCulloch-Pitts neurons, Boltzman machines, and Mean field models, etc.

We have also shown how the analytic stability criteria derived from the Hamiltonian function fares against the stability criterion emerging from the network analysis for a wide range of activation functions, as well as for both additive and shunting networks. The agreement between the two stability criteria is indeed quite close.

For the additive variant, the stability criterion derived from the Hamiltonian has the advantage of predicting onset of Winner-take-all (WTA) behavior in the additive recurrent networks. In fact the two stability criteria (derived from Hamiltonian and network analysis) appear to complement each other. We have also shown how the WTA decision making process be used to predict interesting consequences for studies involving subjective expansion of time in temporal oddball paradigm in psychology.

Furthermore in our previous work [3], we have shown how the Hamiltonian function for the network can be used to derive predictions for psychophysical attributes like reaction times in constructions of biological recurrent networks for humans. We used the Hamiltonian successfully to explain reaction time distributions for visual sense of numbers. To derive the prediction for reaction time (RT) we used only the assumption, *RT* ∝ −*H*, i.e., reaction time should be correlated with the energy that will be needed for resetting the network. Experimentally, the fMRI activation patterns for enumeration and visual working memory task were predicted from the model and subsequently verified through human experimentation [16].

In recent years the understanding of neural codes has provided us with insights that go beyond the concepts of rate coding and it is increasingly more commonplace to speak of temporal codes that use spike timing and phase information in order to transmit and process information reliably [17]–[20]. However, most of these attempts are to look at neural codes after the presentation of stimulus. Recently, some studies have looked at pre-stimulus brain states in MEG and EEG based studies and have found that it is possible to predict conscious detection of stimuli based on pre-stimulus oscillatory brain activity [21]–[24]. For instance in case of near threshold stimuli some researchers have found the prestimulus *α* frequency band modulation to be important [22]. These attempts have drawn a large amount of interest, but have revealed little towards a theoretical or physiological understanding of such phenomena. In section IIB, we have shown the possibility of oscillatory brain states under certain equilibrium conditions for a neural assemble consisting of both feed-forward and recurrent connections. Interestingly, we this leads to an important understanding that the delay between feed-forward and recurrent connections is very important towards understanding the distribution of oscillatory brain states. An alternative approach to oscillatory brain states has been explored in [25].

Overall, the current work not only brings a novel formalism to recurrent neural networks, but also shows way to connect the neural network properties for the purpose of predicting diverse psychophysical parameters in biological systems including but not limited to time perception, enumeration, working memory, oscillatory brain dynamics to name a few.

**Table 1.** Simulation parameters

| Parameter | Value |
| --- | --- |
| $N$ | 10 |
| $\alpha$ | 0.5:0.01:1.5 |
| $\beta$ | 0.3 |
| No. of nodes receiving input | 2 |
| Total duration of simulation in time steps | 2500 |

**Table 2.** Simulation parameters

| Parameter | Value |
| --- | --- |
| $N$ | 10 |
| $\alpha$ | 1.05 |
| $\beta$ | 0.3 |
| No. of nodes receiving input | 2 |
| Total duration of simulation in time steps | 2500 |

## Footnotes

1 In this paper we will ignore the noise terms in the analysis.

2 For example, continuous-time Hopfield networks with network model [8] 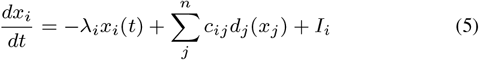 is a special case of Eq. 4 with with *a*_*i*_(*x*_*i*_(*t*)) = 1 and *b*_*i*_(*x*_*i*_(*t*)) = *−λ*_*i*_*x*_*i*_(*t*)+ *I*_*i*_

